# Interspecific introgression reveals a role of male genital morphology during the evolution of reproductive isolation in *Drosophila*

**DOI:** 10.1101/2020.06.03.132100

**Authors:** Stephen R. Frazee, Angelica R. Harper, Mehrnaz Afkhami, Michelle L. Wood, John C. McCrory, John P. Masly

**Author notes:** Author for correspondence: John P. Masly, Department of Biology, University of Oklahoma, 730 Van Vleet Oval, Norman, OK 73019, U.S.A.

## Abstract

Rapid divergence in genital structures among nascent species has been posited to be an early-evolving cause of reproductive isolation, although evidence supporting this idea as a widespread phenomenon remains mixed. Using a collection of interspecific introgression lines between two *Drosophila* species that diverged ∼240,000 years ago, we tested the hypothesis that even modest divergence in genital morphology can result in substantial fitness losses. We studied the reproductive consequences of variation in the male epandrial posterior lobes between *Drosophila mauritiana* and *D. sechellia* and found that divergence in posterior lobe morphology has significant fitness costs on several pre-fertilization and post-copulatory reproductive measures. Males with divergent posterior lobe morphology also significantly reduced the life span of their mates. Interestingly, one of the consequences of genital divergence was decreased oviposition and fertilization, which suggests that a sensory bias for posterior lobe morphology could exist in females, and thus posterior lobe morphology may be the target of cryptic female choice in these species. Our results provide evidence that divergence in genitalia can in fact give rise to substantial reproductive isolation early during species divergence, and they also reveal novel reproductive functions of the external male genitalia in *Drosophila*.

## Introduction

External reproductive structures have long been of interest to evolutionary biologists because of their incredible diversity of form. Among these structures, the external genitalia have attracted particular interest for three primary reasons. First, because external genital structures evolve rapidly among species, they are useful characters in systematics, especially for comparisons among young species (Engel and Kristensen 2013; Kjer et al. 2016). Second, external genitalia provide a powerful model for understanding how sexual selection and sexual conflict affect morphological change over short evolutionary time scales (Eberhard 1985). Third, because of their central role in reproduction, it has been hypothesized that mismatch between interacting male and female genital structures has the potential to cause reproductive isolation (RI) among nascent species (Dufour 1844; De Wilde 1964; Eberhard 1992). Although abundant evidence supports that divergence in genital morphology is often a consequence of sexual selection/conflict (Eberhard 1985; Hosken and Stockley 2004; Simmons 2014; Brennan and Prum 2015), the importance of divergence in genital morphology as a cause of RI has been debated (Shapiro and Porter 1989; Masly 2012).

Nonetheless, several recent studies in a variety of taxa support the idea that morphological divergence in external genitalia can indeed cause RI early during the speciation process via both mechanical and sensory incompatibilities. One well-characterized example of mechanical incompatibility between male and female genitalia occurs among several species of *Carabus* (subgenus *Ohomopterous*) ground beetles, where species divergence in male aedeagus morphology causes substantial damage to the female vaginal appendix during copulation, resulting in reduced reproductive output, damage to the aedeagus, and even female mortality (Sota and Kubota 1998; Nagata et al. 2007; Sota and Tanabe 2010; Kyogoku and Sota 2015). Genomic studies also show that the greatest genetic divergence among these species occurs in regions associated with genital morphology (Fujisawa et al. 2019), consistent with divergence in genitalia as the initial cause of RI in this group. Divergence in male genital bristle morphology between *Drosophila yakuba* and *D. santomea* impedes insertion of the aedeagus during mating, significantly reducing insemination success and often causing damage to the female genitalia (Kamimura and Mitsumoto 2012). And, in the damselfly genus *Enallagma*, divergence in species-specific morphology gives rise to both mechanical incompatibilities that reduce copulation success and sensory incompatibilities where females refuse to mate with males that possess divergent genital morphology, resulting in nearly complete RI (Paulson 1974; Barnard et al. 2017).

Despite these and other examples, the relative importance of divergence in genital morphology as a common contributor to the evolution of RI early during speciation remains unclear. Because many recognized species are often separated by multiple RI mechanisms, isolating any potential contribution of divergence in genital morphology to RI can sometimes be difficult as later-evolved incompatibilities could mask the effect of genital mismatch. One particular set of genital structures that have received considerable attention because of their striking morphological differences among young species and potential for understanding the genetic and developmental bases of complex traits are the epandrial posterior lobes (PLs) in *Drosophila*. The PLs are bilaterally symmetrical cuticular projections on either side of the male external genitalia that insert between female abdominal segments VII and VIII during copulation (Robertson 1988; Eberhard and Ramirez 2004; Kamimura and Mitsumoto 2011), which have evolved among the four species of the *D. melanogaster* complex (Jagadeeshan and Singh 2006) and are essential in each species for securing genital coupling during mating (Frazee and Masly 2015; LeVasseur-Viens et al. 2015). Early tests of the contribution of the PLs to RI showed that mismatch among the species gave rise to defects in copulation duration, sperm transfer, and oviposition, prompting the authors to conclude that divergence in genital morphology causes “cryptic” RI among these species (Price et al. 2001). However, it has been difficult to interpret these results as mate discrimination and divergence in seminal fluid proteins (Sfps) among these species could affect many of these reproductive measures. Two later studies tested the effects of variation in PL morphology on reproductive success by modifying PL size and shape within species, with somewhat contrasting results. In *D. melanogaster*, reductions in PL size and length:width gave rise to decreased copulation duration, reduced sperm transfer, and reduced oviposition, even under competitive fertilization conditions (Frazee and Masly 2015). However, in *D. simulans*, reductions in PL size and modifications in shape showed no apparent effect on copulation duration or sperm transfer, although variation in PL morphology had an effect on male copulation success in a competitive mating environment (LeVasseur-Viens et al. 2015).

A robust test of divergence in PL morphology as a cause of RI requires the generation of species-specific variation in PL morphology in the absence of other RI barriers that separate species. Here, we use an interspecific introgression approach to test the hypothesis that divergence in PL morphology can give rise to substantial incompatibilities at the earliest stages of species divergence. Our test takes advantage of several *D. mauritiana-D. sechellia* genetic introgression lines that possess small chromosomal segments (∼1.5 Mb on average) of the *D. mauritiana* genome within a predominantly *D. sechellia white* (*w*) genomic background (Masly and Presgraves 2007). Pure species *D. mauritiana* possesses small finger-shaped PLs, whereas *D. sechellia* possesses much larger goose-headed-shaped PLs, with a long neck and characteristic “beak.” Several of these *D. mauritiana-D. sechellia* introgression lines possess interspecific variation in male PL morphology including transgressive variation in PL size, whereas others possess morphology that is similar to *D. sechellia w* (Masly et al. 2011). Importantly, these introgression lines do not possess any strong RI barriers that are observed between the two pure species, such as intrinsic hybrid sterility or behavioral isolation (Masly and Presgraves 2007; Cattani and Presgraves 2009; Masly et al. 2011; McNabney 2012). We use these lines in mating experiments to *D. sechellia w* females and quantify several reproductive measures to identify the potential effect(s) of divergent PL morphology on fitness loss.

## Material and Methods

### *Drosophila* stocks

*Drosophila* stocks were reared on cornmeal-molasses-agar medium at 25°C and 65-70% relative humidity under 12-hour light:dark conditions. The *D. mauritiana-D. sechellia* introgression lines used in our study represent the full range of PL morphologies observed among these lines (Fig. 1) and include lines that broadly possess significant reductions in PL size compared to *D. sechellia w* (*Q1(A)* and *4C2(A)*), lines that possess significant differences in shape compared to *D. sechellia w* (*3Q1(A), DEE1(B), I1(B), NENEH2(A)*), lines that possess larger size, but similar shape compared to *D. sechellia w* (*4G4C(A)*), and lines that possesses both larger size and a difference in shape (*2U1(C)* and *2H3(B)*). We also included two “introgression control” lines in our study (*YAR1(A)* and *4G5(A)*) that possess PL morphology that is not significantly different from that of *D. sechellia w*. This collection of *D. mauritiana-D. sechellia* introgression lines also mirrors those used in a previous study that quantified PL insertion-site wounds suffered by females during mating with males that possess interspecific PL morphologies (Masly and Kamimura 2014).

**Figure 1.**
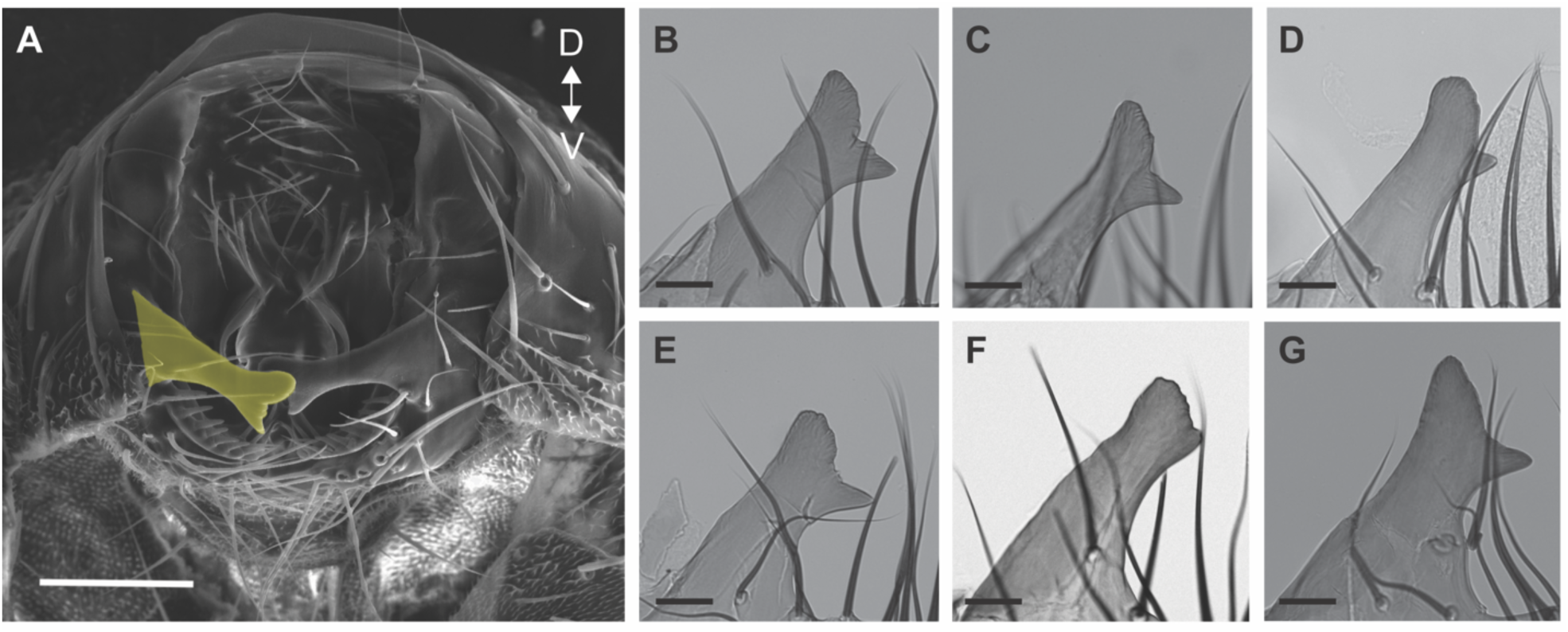
Examples of epandrial posterior lobe morphological variation among genotypes. (A) Male terminalia in *D. sechellia*. One of the posterior lobes is shaded yellow. D and V indicate the dorsal and ventral axes. (B) *D. sechellia w*; (C) *Q1(A)*, an introgression genotype that possesses significantly smaller PL size compared to *D. sechellia w*; (D) *3Q1(A)*, an introgression genotype that possesses significantly different shape compared to *D. sechellia w*; (E) *YAR1(A)*, an introgression control genotype with PL morphology similar to *D. sechellia w*; (F) *DEE1(B)*, an introgression genotype that possesses significantly different shape compared to *D. sechellia w*; (G) *4G4C(A)*, an introgression genotype that possess larger size, but similar shape compared to *D. sechellia w*. Scale bars: (A) 100 μm, (B-G) 25 μm.

### Mating assays

Three-day old virgin *D. sechellia w* females were placed in eight-dram food vials with one to five three-day old virgin males of a particular genotype within one hour of first light. Once copulation occurred, all males that were not copulating were immediately removed from the mating vial via aspiration. For each successfully copulating pair, we recorded copulation duration (minutes) and the copulation orientation of the male during mating. Copulation orientation was scored as abnormal if a male maintained an abnormal mounting position (skewed at an angle of at least 45 degrees to either side of the female or leaning straight back at a 90-degree angle) for at least one continuous minute during the entire copulation period. Males and females were immediately separated after copulation ended and females were frozen immediately to enable quantification of male sperm transfer to the reproductive tract. We dissected the female reproductive tract in 1X PBS on a glass slide and removed the spermathecae, seminal receptacle, and uterus/common oviduct. The contents of these organs were then spread on the slide, allowed to dry, fixed in 3:1 methanol:acetic acid, and stained with 0.2 μg/ml DAPI to visualize sperm nuclei. Sperm nuclei were quantified using 100X magnification. We scored sperm number twice for all samples with consistent results (*r*=0.98).

Individual males were isolated for three days following their initial mating to replenish expended sperm before being mated individually with a new *D. sechellia w* virgin female. Mated females were transferred to a new food vial every 3 days for 15 days. We recorded the number of eggs that were laid, number of eggs that hatched, and the total number of progeny that emerged from each of the five vials. Progeny were scored up to day 19 after the adults were first introduced into each new vial. We tested an average of n=30 males for each genotype we studied, and each set of mating experiments was scored blind with respect to male genotype.

### Survival assays

Three-day old virgin *D. sechellia w* females and virgin males were paired individually within an hour of first light in food vials and observed to mate. Mated females remained isolated in individual food vials and were observed daily to record mortality. Surviving females were transferred to a fresh food vial every five days until all females had died. Survivorship was recorded as the number of days a female survived after mating. We tested an average of n=30 females in matings with males from each of the genotypes used in our study.

### Effect of artificial wounds on egg laying

To artificially produce wounds at the PL insertion sites in virgin females, we collected newly-eclosed *D. sechellia w* females and anesthetized them under light CO_2_. We then gently inserted an unsterilized 0.25 mm diameter insect pin (Bioquip Products) between abdominal segments VII and VIII on either side of the abdomen at the site of PL insertion during copulation. This sized insect pin substantially exceeds the size of the *D. sechellia w* PL, and wounds were evident by trace amounts of hemolymph that leaked out at the insertion sites. These females were allowed to recover for four days in isolation before being placed in individual food vials and transferred to a new vial every three days for nine days. Control four-day old virgin *D. sechellia w* females that were not wounded were likewise placed in food vials and transferred. We recorded the total number of eggs laid across all three vials. To assay artificial wounds in mated females, virgin males and females were collected and aged in isolation for three days. After this time, one virgin male and one virgin female were paired together in a food vial, and the males were removed after 24 hours. Females were then lightly anesthetized using light CO_2_ and wounded with an insect pin as described above. Wounded experimental females and unwounded control females were returned to individual food vials and transferred to a new food vial every three days for nine days.

### Enzyme-linked immunosorbent assays (ELISAs)

To estimate seminal fluid protein (Sfp) transfer from a single mating, we performed ELISAs using an antibody against Sex Peptide (SP) following the protocol described in ref. (Sirot et al. 2009). Three-day old virgin *D. sechellia w* females and experimental and control males were mated individually as described above and copulation duration was recorded for each successful mating. Immediately after mating, males and females were separated and flash frozen in liquid nitrogen. Samples were stored at −80°C until dissection.

We generated SP standards by dissecting the accessory glands from 30 virgin *D. sechellia w* males and homogenizing them in a microcentrifuge tube containing 60μl of 10% Dulbecco’s phosphate buffered saline (DPBS; 14 mM NaCl; 0.2 mM KCl; 0.1 mM KH_2_PO_4_; 0.7 mM Na_2_HPO_4_) with cOmplete™ Protease Inhibitor (PI) Cocktail Tablets (Roche). Accessory glands were homogenized for 30 sec., then the pestle was then rinsed with 1.2ml of 10% DPBS with PI. Two hundred microliters of the homogenate were serially diluted (dilution series: 1, 1/2, 1/4, 1/8, 1/16, 1/32, 1/64, 1/128, 1/256, 1/512) and 50μl of each dilution was added to Immulon™ 2 HB flat bottom 96-well ELISA plates (Thermo Scientific) in triplicate. We also included 10% DPBS with PI on each plate in triplicate as a blank for the absorbance measurements.

The uterus from each mated *D. sechellia w* female was dissected in ice-cold 10% DPBS with PI and placed into a microcentrifuge tube containing 20μl of 10% DPBS with PI. Each uterus was homogenized for 30 sec., and the pestle was then rinsed with 200μL of 10% DPBS with PI. Each of the samples was then serially diluted (dilution series: 1, 1/2, 1/4, 1/8, 1/16) and 50μl of each dilution was added to the plate. Once filled, plates were sealed and placed on an orbital shaker overnight at 4°C. The liquid was then aspirated out and the bound sample in each well was incubated in 100μl of blocking buffer (5% nonfat milk, 0.05% Tween-20 in 1X DPBS) on an orbital shaker for 1 hr. at room temperature (RT) followed by 50μl of rabbit anti-SP (1:750 dilution in blocking buffer) for 2 hrs. at RT. The SP antibody was removed, and each well was washed three times with 0.05% Tween-20 in 1X DPBS. Samples were then incubated with 50μl goat anti-rabbit horseradish peroxidase (1:2,000 in blocking buffer) for 1 hr. at RT then washed as before. Following these washes, 100μl of 3,3’, 5,5’-tetramethylbenzidine substrate was added to each well and incubated for 15 min. at RT. Each reaction was quenched with 100μl 1M HCl, and the absorbance of the wells was immediately measured at 450nm (OD_450_) using an EL 800 Universal Microplate Reader (Bio-Tek Instruments).

To generate the standard curves for each plate, the average OD_450_ of the blank was subtracted from the average OD_450_ of each dilution factor, and these values were plotted against the dilution factor OD_450_ to obtain a linear equation with *R*^2^ values for each plate (*R*^2^ values among plates were 0.98-0.99). To enable comparisons across all plates, we used a linear conversion to standardize OD_450_ values, so that the standard curves each had a slope of one and a *y*-intercept of zero. We report the results using standardized OD_450_ values from our dilution factor of 1/4 treatments, but our analyses using the OD_450_ values from other dilutions yield similar results.

### Morphological measurements

Left and right PLs and epandrial ventral plates (lateral plates) were dissected from males, mounted in polyvinyl alcohol medium (Bioquip Products) on glass slides, and imaged at 200X magnification. The outline of each PL was manually traced using ImageJ (Rasband 1997-2019) and enclosed with an artificial baseline drawn in line with the lateral plate. Each closed contour was then converted into (*x,y*) coordinates that were used in elliptical Fourier analysis (Kuhl and Giardina 1982; Ferson et al. 1985), which allows comparison of disparate shapes with high precision (Kuhl and Giardina 1982; Lestrel 1997) and effectively captures morphological variation in the PL both between and within species (Liu et al. 1996; Macdonald and Goldstein 1999; Zeng et al. 2000; Masly et al. 2011; McNeil et al. 2011; Masly and Kamimura 2014; Frazee and Masly 2015; Takahara and Takahashi 2015; Takahashi et al. 2018; Tanaka et al. 2018). For each PL we obtained 80 Fourier coefficients and used principal components analysis (PCA) to reduce the number of variables that describe variation in PL morphology. Elliptical Fourier coefficients were adjusted to standardize location, orientation, and handedness within the coordinate plane prior to PCA. We selected one PL at random from each individual we dissected to include in our PCA, and PCA was performed using singular value decomposition of the elliptical Fourier coefficient data matrix. The first three PC scores explained approximately 75 percent of the morphological variation in our dataset and were used to represent PL morphology in statistical analyses. Although it is difficult to assign exact morphological correlates to each PC score, in general PC1 correlates with PL area, PC2 correlates with PL length:width, and PC3 with prominence of the characteristic *D. sechellia* “beak” structure (Fig. 1). The length from the tibiotarsal joint to the tibiofemoral joint of the male forelegs was measured to provide an estimate of overall body size (Catchpole 1994; Kacmarczyk and Craddock 2000; Siomava et al. 2016).

### Statistical analyses

The effect of variation in morphology on pre-fertilization reproductive measures was tested using multivariate analysis of variance (MANOVA) with the reproductive measures as the response variables and the representations of PL morphology plus tibia length as explanatory variables. Tests for effects among reproductive measures, the effect of pre-fertilization measures, PL morphology, and tibia length on post-copulatory reproductive measures, and tests for the effect of PL morphology, tibia length, and copulation duration on Sfp transfer were all performed using analysis of variance.

Copulation orientation was modeled as a binary response variable and analyzed using a GLM with PL morphology, tibia length, and copulation duration as explanatory variables. Egg hatch success was modeled as a proportion, and a GLM was used to test the effect of PL morphology, tibia length, and copulation duration on egg hatch success. Because these data were overdispersed, we corrected for overdispersion by fitting the model using quasibinomial distributed errors with a logit link function. Female survivorship data was analyzed using a Cox proportional hazard model with mortality as a constant hazard.

We used all of our available observations to maximize our sample size for each statistical test that we performed. All statistical analyses were performed using R release 3.5.3 (R Core Team 2019). Figures were constructed using either the base graphics package in R or the package ggplot2 (Wickham 2009). Means are reported ±1 s.e.m. Post-hoc tests were performed using the Tukey method.

## Results

### Posterior lobe morphology affects multiple reproductive fitness measures prior to fertilization

Although *D. mauritiana-D. sechellia* introgression males do not display any behavioral isolation that prevents mating (McNabney 2012), we observed that copulation latency can often be prolonged with individual mating pairs in food vials. This was observed even for individual male-female mating pairs of *D. sechellia w* placed in food vials. To facilitate copulation in a reasonable observation period, we included additional males in matings with a single *D. sechellia w* female. In contrast to what has been observed in *D. melanogaster* (Bretman et al. 2013), the number of males in a vial had no effect on either pre-fertilization measures (copulation latency, copulation duration, copulation position, sperm transfer; MANOVA, *F*_16,923_=1.39, *P*=0.14) or post-copulatory measures (total oviposition, total offspring; *F*_8,726_=1.39, *P*=0.31). We thus performed our statistical analyses without including the number of males per vial as a covariate.

We tested the effect of male morphology on the four pre-fertilization phenotypes that we measured. Although tibia length showed a significant effect on copulation latency with larger males exhibiting shorter latencies (MANOVA; *F*_*1*,299_=20.3, *P*=9.4 x 10^−6^), tibia length had no effect on copulation duration (*F*_1,299_=2.01, *P*=0.16), copulation positioning (*F*_1,299_=1.96, *P*=0.16), or sperm transfer to the female (*F*_1,299_=0.31, *P*=0.56). In contrast, PL morphology had significant effects on all four reproductive measures (copulation latency: *F*_3,299_=8.15, *P*=3.1 x 10^−5^, copulation duration: *F*_3,299_=5.64, *P*=9.0 x 10^−4^, copulation positioning: *F*_3,299_=4.55, *P*=0.003, sperm transfer: *F*_3,299_=3.56, *P*=0.015). Because three pre-fertilization traits that we measured culminate in sperm transfer to the female, it is possible that copulation latency, copulation duration, and/or copulation positioning may affect levels of sperm transfer from a single mating. We thus tested the effects of these three measures on sperm transfer and found that none had a significant effect (copulation latency: *F*_1,305_=0.46, *P*=0.5, copulation duration: *F*_1,305_=0.82, *P*=0.37, copulation positioning: *F*_1,311_=2.19, *P*=0.14). Intrinsic deficits in male sperm abundance and motility also do not explain the reduced sperm transfer amounts, as we observed no significant differences among genotypes (*χ*^2^=13.3, df=22, *P*=0.92; Supporting information; Table S1).

The most visually striking mating trait during our observations was male orientation on the female during the duration of copulation. Males of certain *D. mauritiana-D. sechellia* introgression lines would often experience difficulty maintaining a normal copulation position on the back of the female during mating. In particular, these males would maintain copula skewed at an angle of 45 degrees to either side of the female or lean straight back at a 90-degree angle. We modeled copulation position as a binary trait (normal vs. abnormal) and tested the effects of PL morphology, tibia length, and copulation duration on male positioning using a generalized linear model (GLM). We found that although tibia length (*P*=0.15) and copulation duration (*P*=0.07) had no effect on male positioning, PL morphology had a significant effect on a male’s ability to maintain the proper orientation (*P*=0.01). In particular, males with smaller PLs were more often unable to maintain copulation orientation (Fig. S1). Taken together, the results of our analyses show that males with smaller or abnormally-shaped PLs remained in copula for longer periods, suffered abnormal copulation positioning more frequently, and transferred fewer sperm than males that possessed PL morphology that was either similar to *D. sechellia w* or larger than *D. sechellia w*.

### Posterior lobe morphology affects female oviposition and contributes to fertilization success

In *D. melanogaster*, females that mate with males possessing smaller or narrower PLs significantly reduce the number of eggs that they lay from a single mating (Frazee and Masly 2015). We found a similar effect of PL morphology when *D. mauritiana-D. sechellia* introgression males mated with *D. sechellia w* females (*F*_3,290_=8.97, *P*=1.1 x 10^−5^), although there was no effect of tibia length (*F*_1,290_=1.06, *P*=0.30), copulation duration (*F*_1,290_=1.04, *P*=0.31), copulation positioning (*F*_1,296_=0.40, *P*=0.53), or sperm transfer (*F*_1,305_=1.19, *P*=0.28) on oviposition amounts. Also similar to what was observed within *D. melanogaster*, females that mated with males possessing smaller PLs laid fewer eggs than those mated to males with larger PLs (Fig. 2; *z*=2.32, *P*=0.02).

**Figure 2.**
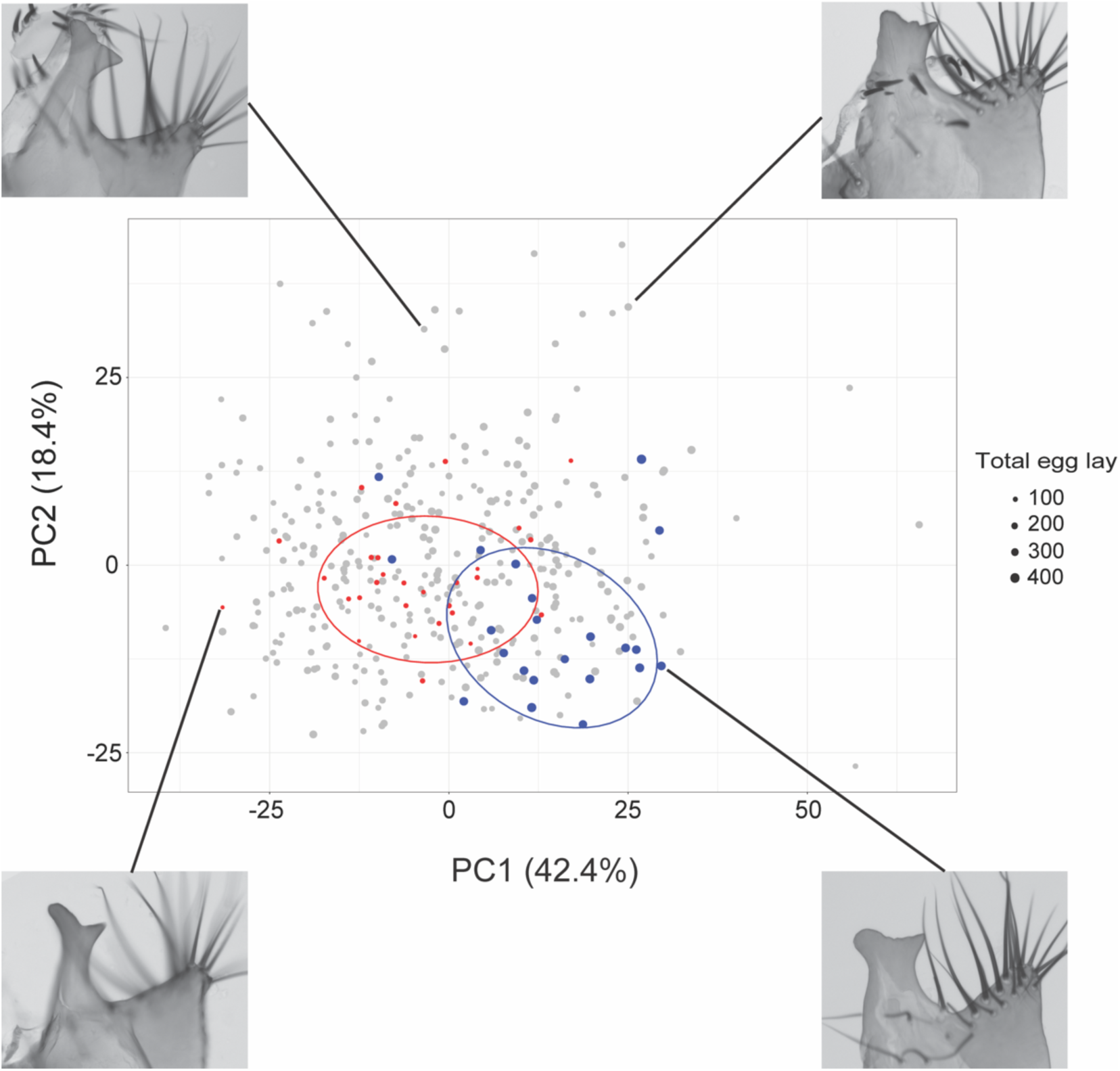
Variation in posterior lobe morphology affects oviposition. Variation in posterior lobe morphology is shown across the distribution of principal component 1 (PC1) and principal component 2 (PC2). The number of eggs oviposited by females after mating is shown by the size of each plotted point. Oviposition amounts in the lowest and highest tenth percentiles are shown in red and blue, respectively, with 75% normal-probability ellipses. Images of posterior lobes show representative examples of the distribution in morphology across the PC1-PC2 axes. Numbers in parentheses show the proportion of morphological variation explained by each principal component.

There was high correlation between the number of hatched eggs and the number of offspring across genotypes (*r*=0.86), consistent with the lack of substantial viability effects observed in heterozygous introgression males (Masly and Presgraves 2007; Cattani and Presgraves 2009). We thus used the ratio of hatched eggs to total eggs laid as an estimate of fertilization success. Our tests revealed that PL morphology (GLM; *P*=8.4 x 10^−4^), tibia length (*P*=0.017), and copulation duration (*P*=0.028) all had significant effects on egg hatch, but copulation position (*P*=0.18) and sperm transfer amount (*P*=0.12) did not. The aspect of PL morphology that had the greatest effect on egg hatch was PC2 (*t*=2.75, *P*=0.004), which roughly corresponds to PL length:width (Fig. 2).

### Variation in oviposition is not a consequence of reduced seminal fluid protein transfer or posterior lobe wounding

Introgression males that possess smaller or misshapen PLs transfer fewer sperm than pure species *D. sechellia w* males or males that possess larger PLs. Several Sfps are known to affect oviposition in *Drosophila* (Wolfner 1997; Chapman and Davies 2004), thus the possibility exists that in addition to transferring fewer sperm in a single mating, these introgression males might also transfer less Sfps, which could result in the observed reduction in egg laying. To estimate Sfp transfer amounts, we performed ELISAs to quantify the amount of SP transferred to the female reproductive tract from a single mating. SP is a major component of the male ejaculate and is functionally conserved within the *D. melanogaster* species group (Tsuda et al. 2015; Tsuda and Aigaki 2016). Although the introgression lines differ in the amount of SP they transfer, there was no significant effect of copulation duration (*F*_1,49_=0.213, *P*=0.65), tibia length (*F*_1,49_=2.748, *P*=0.10), or PL morphology (*F*_3,49_=0.41, *P*=0.75) on SP transfer amount during mating (Fig. 3). Interestingly, males from one of the introgression lines that possess the smallest PLs transfer the largest amounts of SP (*4C2(A)*, Fig. 3D), counter to the expectation that increasing amounts of Sfps might give rise to increased egg laying. Thus, it does not appear that reduced Sfp transfer explains the reduced oviposition in matings to introgression males with smaller or abnormally-shaped PLs.

**Figure 3.**
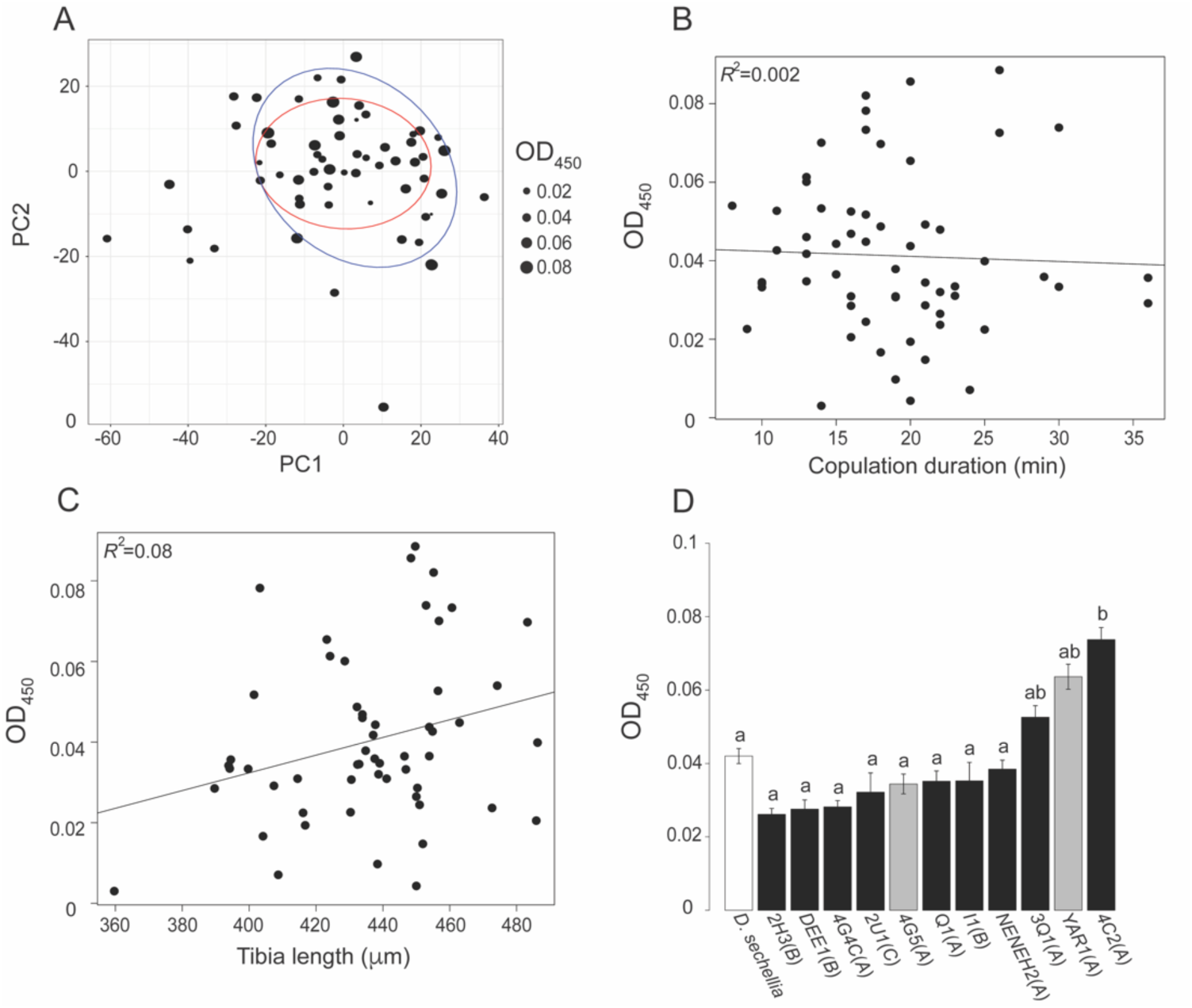
Posterior lobe morphology has no effect on Sex Peptide transfer during mating. (A) Variation in posterior lobe morphology is shown across the distribution of principal component 1 (PC1) and principal component 2 (PC2). SP amount transferred to females after mating is shown by the size of each plotted point. Red and blue 75% normal-probability ellipses show the SP amounts in the lowest and highest tenth percentiles, respectively. (B) Correlation between SP abundance in the female reproductive tract after a single mating and copulation duration and (C) tibia length. (D) Average SP transfer amounts among genotypes. White shows *D. sechellia w*, black bars show *D. mauritiana-D. sechellia* introgression lines with divergent posterior lobe morphologies, and grey bars show *D. mauritiana-D. sechellia* introgression lines with *D. sechellia*-like posterior lobe morphology. Statistically homogeneous groups were assigned using **α**=0.05.

Oviposition could also be reduced as a consequence of species-specific divergence in Sfps. Sfps diverge rapidly among *Drosophila* species (Panhuis Tami et al. 2006), thus any substantial protein sequence divergence in Sfps encoded by *D. mauritiana* alleles within the introgression regions could be incompatible with their interacting partners in the female *D. sechellia* reproductive tract. We identified genes within each *D. mauritiana* introgression that encode Sfps that are transferred to the female during mating among species of the *D. melanogaster* subgroup (Findlay et al. 2008; Findlay et al. 2009; Sepil et al. 2019) and checked their molecular evolutionary rates using the available population genomic data from comparisons between *D. simulans* and *D. mauritiana* (Garrigan et al. 2012) and *D. melanogaster* and *D. simulans* (Begun et al. 2007). McDonald-Kreitman test results show that none of the 13 transferred Sfps that exist within the introgression regions are evolving by positive natural selection (Table S2). We also examined evolutionary rates for the known sperm proteins in *D. melanogaster* (Dorus et al. 2006; Wasbrough et al. 2010) that are encoded by genes within the *D. mauritiana* introgressions. Although some of these genes show a signature of positive selection (Table S2), it is unclear from their known or predicted functions whether these proteins localize to the sperm cell membrane where they could potentially interact directly with the female reproductive tract. Moreover, the transfer of sperm alone to the female has a negligible effect on oviposition compared to the effect of Sfps (Heifetz et al. 2001), thus it seems unlikely that incompatible interactions with divergent sperm proteins would give rise to such significant reductions in oviposition that we observed.

Because introgression males with divergent PL morphology cause wounds at the PL insertion sites more often than *D. sechellia w* males (Masly and Kamimura 2014), it is possible that the reduced oviposition we observed is a consequence of mated females diverting resources from reproduction to immunity. To test this idea, we used fine insect pins to generate artificial wounds at each PL insertion site on both virgin and inseminated *D. sechellia w* females and compared oviposition rates between wounded and unwounded individuals. Interestingly, wounded virgin females laid slightly more eggs than unwounded virgin females (32 ± 8; n=12 vs. 28 ± 5; n=17), although this difference was not significant (*t*_27_=0.47; *P*=0.32). Inseminated females that were wounded artificially also laid slightly more eggs than inseminated females that were not wounded artificially (63 ± 5; n=16 vs. 59 ± 6; n=17), although this difference, too, was not significant (*t*_31_=-0.56; *P*=0.58). Thus, our results show that the reduced oviposition in mates of males with smaller or misshapen PLs does not appear to be a consequence of either Sfp transfer amount or divergence, nor resource reallocation as a consequence of wounds suffered during mating.

### Females mated to males with divergent posterior lobe morphologies suffer decreased longevity

Because males with divergent PL morphologies wound females more severely than either *D. sechellia w* males or males with larger than normal PLs (Masly and Kamimura 2014), it is possible that these males might also reduce female lifespan and further reduce female fecundity, similar to the deleterious effects of divergent genital morphology observed in some interspecific crosses (Masly 2012). We quantified *D. sechellia* female longevity after a single mating and found that longevity among females mated with males of different genotypes is significantly different (*χ*^2^=140.1; df=11; *P*<2.2 x 10^−16^; Fig. S2). In particular, the *D. mauritiana-D. sechellia* introgression males that wound significantly more than *D. sechellia w* males caused earlier female mortality (matings with introgression males: 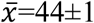 days; matings with *D. sechellia w* males: 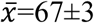 days; *χ*^2^=49.5, df=2, *P*<1.84 x 10^−11^; Fig. 4). Interestingly, females that mated with introgression males of genotypes that do not wound significantly more than *D. sechellia w* (including two genotypes that possess divergent PL morphologies) also experienced significantly earlier mortality compared to those mated with *D. sechellia w* males (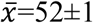 days; *P*=3.2 x 10^−4^, Fig. 4). Although we cannot completely exclude the possibility that Sfps from the *D. mauritiana-D. sechellia* introgression males have slightly deleterious effects on *D. sechellia w* female life span (*e.g.*, Chapman et al. 1995; Holland and Rice 1999), it is worth noting that although these introgression males do not wound significantly more than *D. sechellia w* males statistically, almost all of these genotypes wound females more frequently than *D. sechellia w* (Masly and Kamimura 2014). The one exception was an introgression control genotype (*4G5(A)*) that wounds females less than *D. sechellia w* males (Masly and Kamimura 2014), and shows longer female longevity after mating compared to *D. sechellia w* (Fig. 4), although this difference is not significant (*χ*^2^=1.42, df=1, *P*=0.23).

**Figure 4.**
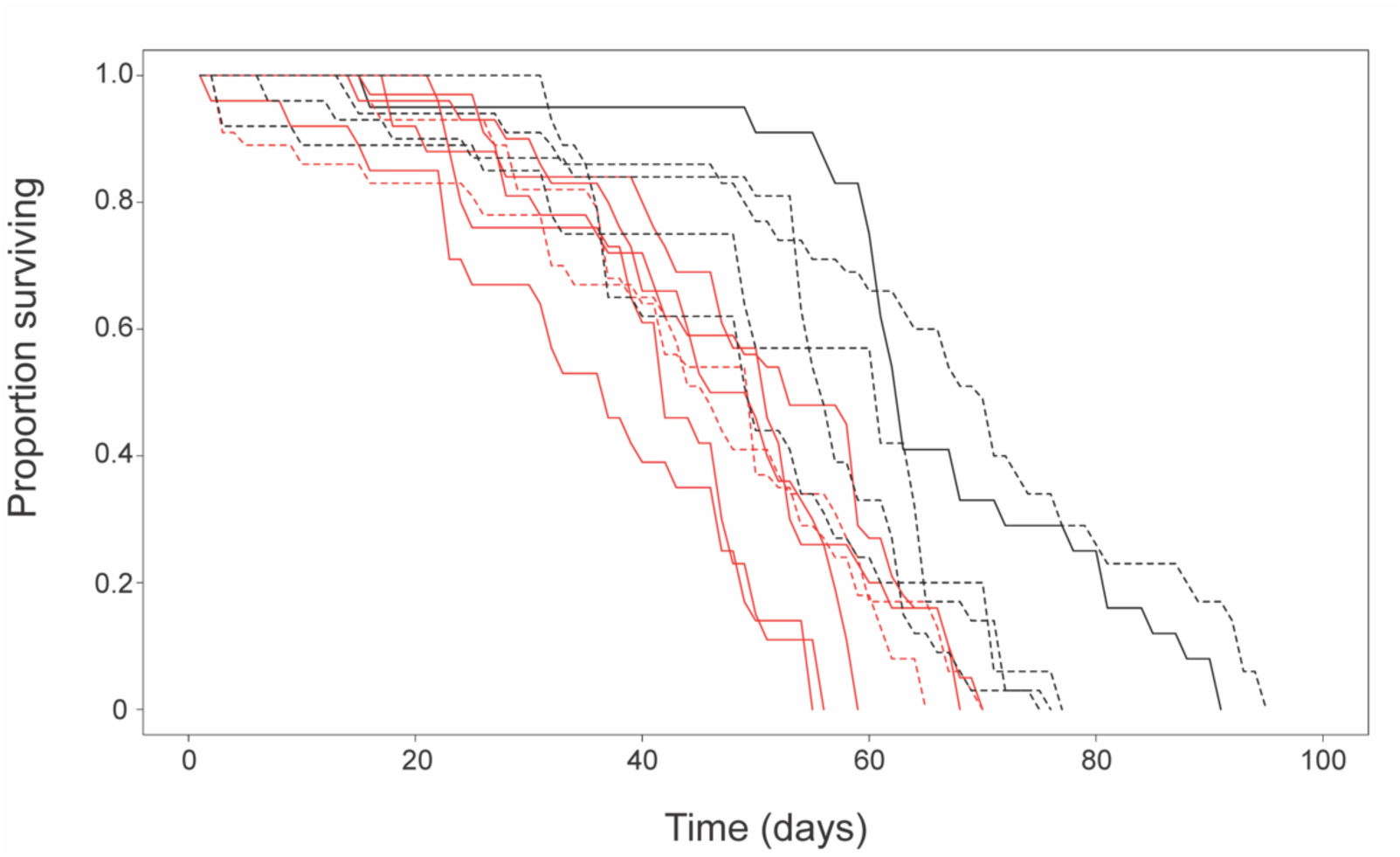
Divergent posterior lobe morphology causes earlier female mortality post-mating. Survivorship curves for females that mate with *D. sechellia w* males (solid black line), *D. mauritiana-D. sechellia* introgression males that wound females significantly more than *D. sechellia w* (solid red line), introgression males that possess divergent posterior lobe morphologies, but do not wound females significantly more than *D. sechellia w* (dashed red line), and introgression males that possess *D. sechellia*-like posterior lobe morphology and do not wound females significantly more than *D. sechellia w* (dashed black line).

## Discussion

Our results show that even modest divergence in PL morphology can significantly decrease fitness, and thus contribute to the evolution of RI. Although divergence in PL morphology did not cause complete RI among the genotypes we studied, the fitness deficits suffered by both sexes provides proof-of-principle support that mismatched genitalia can contribute to RI early during speciation by providing substantial selective pressure on reinforcement (*e.g.*, Comeault and Matute 2016). Previous studies in *D. simulans* have shown that the PLs serve an important function for copulation success in a competitive mating environment (LeVasseur-Viens et al. 2015), and together with the present results and those within *D. melanogaster* (Frazee and Masly 2015), these data suggest that PL morphology alone could potentially give rise to strong RI at early stages of species divergence in the *D. melanogaster* complex.

Similar to the consequences of variation in PL morphology within *D. melanogaster*, our results show that interspecific variation in PL morphology among the *D. mauritiana-D. sechellia* introgression lines affects several pre-fertilization and post-copulatory reproductive measures, and they are also generally consistent with those obtained from crosses among pure species within the *D. simulans* clade (Price et al. 2001). In particular, we found that divergence in PL morphology can cause deleterious fitness consequences on sperm transfer, oviposition, and egg hatch. Our data also show that post-copulatory fitness deficits do not appear to be a due to divergence in Sfps between species. Notably, we found that the direction of the reproductive consequences with respect to PL morphology was similar between our study and the study comparing crosses among the pure species. Specifically, when pure species females mate with males possessing smaller PLs compared to those of conspecifics, oviposition and egg hatch success are both reduced. Conversely, increases in PL size beyond that which is typical of conspecific males often gives rise to increases in copulation duration and sperm transfer amounts. When this PL size increase is modest, there appears to be little effect on fitness in single matings, although in the case of substantial increases (*e.g., D. simulans* male x *D. mauritiana* female) sperm transfer can be so voluminous that the sperm mass obstructs the passage of eggs (Price et al. 2001).

Unlike the results of crosses among the pure species, we found that males possessing divergent PL morphology decrease the longevity of their mates. These differing results might be explained by variation in the severity of wounds induced by male external genital structures during mating. Males of all four species of the *D. melanogaster* complex cause wounds during mating (Kamimura and Mitsumoto 2011), and a previous study using the *D. mauritiana-D. sechellia* introgression lines showed that reductions in PL size or abnormal PL shape increased the frequency of wounding to *D. sechellia w* females, whereas increases in PL size had no effect on wounding compared to controls (Masly and Kamimura 2014). Crosses between pure species could also vary in their degree of wounding, although this has not been measured. But, if the reduction in female longevity we observed is a consequence of copulatory wounding, then some interspecific crosses might not produce the same severity of wounds that is observed among the *D. mauritiana-D. sechellia* introgression lines. The study among pure species also measured the effects of mating on longevity within and between *D. simulans* and *D. mauritiana*, so another possible explanation for the differing longevity results is that *D. sechellia* females could be more sensitive to mating wounds compared to its sister species.

Although PL morphology had a significant effect on sperm transfer amounts, we found that it appears to have little effect overall on transfer of Sfp amount during mating. However, our current data do not allow us to identify whether PL morphology has a direct effect on oviposition and fertilization for two reasons. First, because SP associates with the sperm tail (Peng et al. 2005) and affects release of sperm from the female’s storage organs (Avila et al. 2010) it is possible that females mated to males who transfer fewer sperm during mating, store fewer sperm and consequently store lesser amounts of Sfps like SP. The long term (*e.g.*, beyond one or two days) deficit of Sfp titers could potentially have consequences on oviposition and fertilization several days after mating. Our data show that there was no significant effect of initial sperm transfer amount on oviposition and egg hatch, and the amount of sperm transferred initially exceeds what is typically stored by females in this species group (Fowler 1973; Manier et al. 2010). Thus, it seems reasonable that variation in sperm storage is not the ultimate cause of the observed reductions in oviposition and fertilization. Second, although our data show that the amount of SP transferred during mating is fairly uniform across genotypes, we cannot exclude the possibility that the relative proportions of other Sfps transferred to the female differ across genotypes, and this could potentially affect oviposition rates. Despite these considerations, our results support a significant contribution of PL morphology (either directly or indirectly) to variation in oviposition and fertilization in *Drosophila*.

External genitalia evolve rapidly compared to other morphological structures, and this pattern is widespread among taxa with internal fertilization (Eberhard 1985). Considering the fitness effects of genital mismatch that we observed here, divergence in genital morphology might prove to be a key event during the early stages of speciation among many species. The results of our study also complement a growing body of work that clearly demonstrates that mismatch in reproductive structures can give rise to substantial reproductive incompatibilities. One recent study using *D. mauritiana-D. simulans* introgression lines generated morphological modifications in multiple male terminal structures, which caused severe mechanical incompatibilities that resulted in copulation and insemination defects (Tanaka et al. 2018). We found that divergence in even a single genital structure can cause mechanical incompatibilities, and our results also suggest that the PLs in *Drosophila* might function in a sensory capacity that affects the female reproductive processes of oviposition and fertilization. In particular, our data provide evidence that the PLs function in cryptic female choice, whereby a female might reduce her oviposition and fertilization rates effectively limiting her level of reproductive investment from “less attractive” males (Eberhard 1996). The neural circuits by which *Drosophila* females respond to tactile mating stimuli are beginning to be uncovered (Shao et al. 2019), which promises to reveal avenues for future inroads to understanding the mechanistic bases of how sexual selection shapes phenotypic evolution that is important for male-female mating interactions.

## Acknowledgements

We thank R. Knapp, I. Schlupp, and L. Weider for helpful advice during the course of this project and C. Elenwo for technical help. We also thank M. Wolfner for generously sharing the SP antibody, and D. Presgraves and C. Muirhead for providing the *D. mauritiana-D. simulans* McDonald-Kreitman test results for the Sfp genes. M. Wolfner provided helpful comments on an earlier version of this manuscript. The research reported in this publication was supported by funds from NSF CAREER Award IOS 1453642 to JPM. The content is solely the responsibility of the authors and does not necessarily represent the official views of the National Science Foundation or the University of Oklahoma.

## Author contributions

JPM conceived of the project; SRF, ARH, MA, MLW, JCM, and JPM performed the experiments and collected the data; SRF, ARH, MA, and JPM analyzed the data; JPM wrote the manuscript with input from the coauthors. All authors read and approved the final manuscript.

## Data accessibility

The data described in this paper are deposited in the Dryad Digital Repository, doi:xx.xxxx.

