## Supporting information for "Interspecific introgression reveals a role of male genital morphology during the evolution of reproductive isolation in *Drosophila*"

### Male sperm abundance and motility assay

We measured the abundance and motility of sperm in the testes of males from each of the genotypes we tested in our experiments using the assay described in Orr (1992) and Masly et al. (2006). Virgin males were aged for five to seven days in isolation before their testes were removed, dissected in 1X PBS, and immediately examined under a compound microscope with dark-field optics at 100X magnification. We classified each male into three sperm motility classes: "Many" for males with abundant motile sperm present that fill multiple fields of view; "Few" for males that show only a few localized patches of motile sperm; "None" for males that possess no motile sperm.

**Table S1. Male sperm motility and abundance.**

**Table S2. Molecular evolutionary rates for seminal fluid protein genes and sperm protein genes within the *D. mauritiana-D. sechellia* introgressions.** Yellow-highlighted rows indicate genes encoding seminal fluid proteins that are transferred to the female during mating.

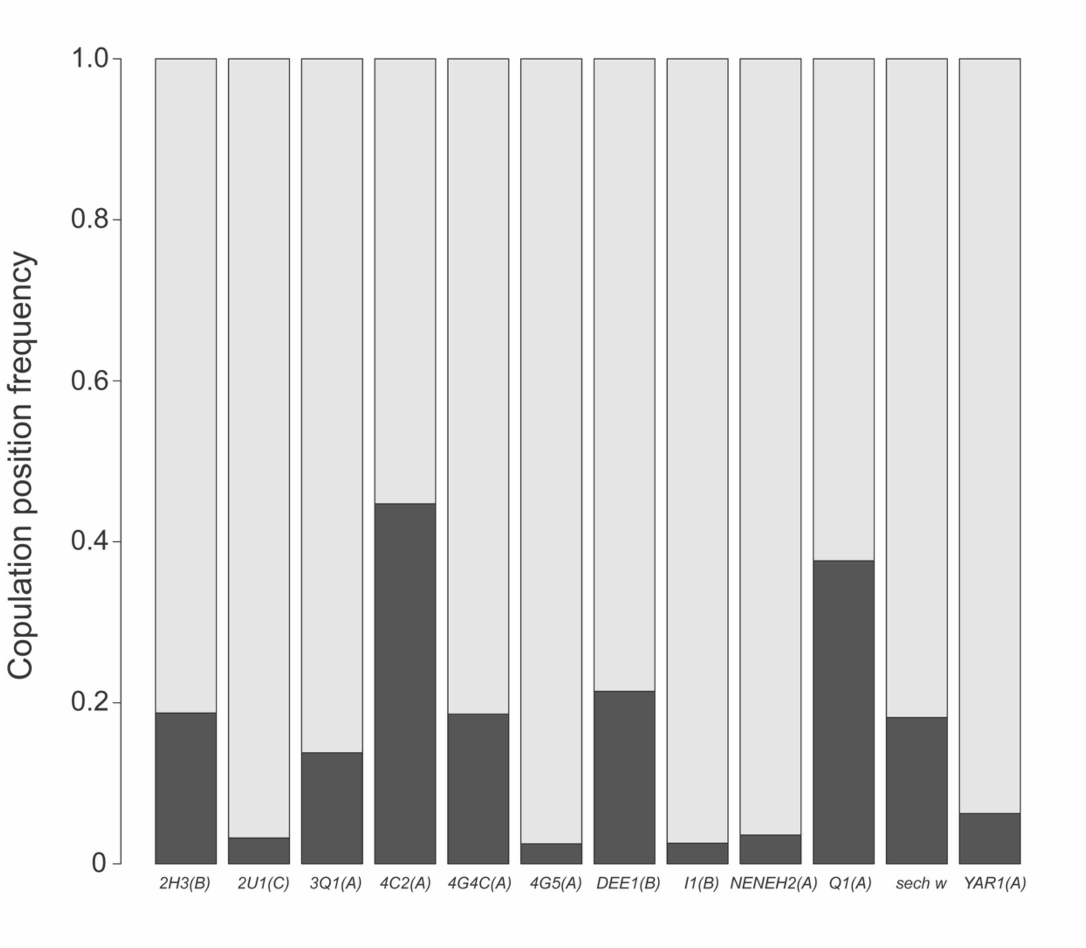


**Figure S1. Copulation positioning differs among genotypes.** Light grey indicates the frequency of normal copulation positioning during mating; black indicates the frequency of abnormal positioning during mating.

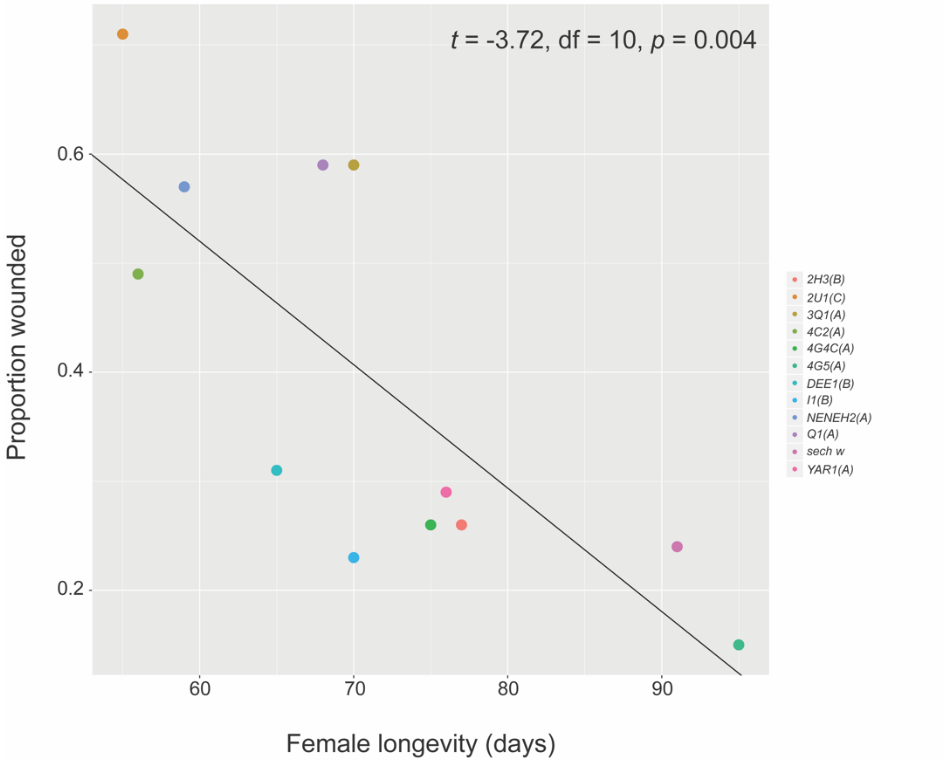


**Figure S2. Effect of posterior lobe wounding on female longevity.**
